# The Real-time fMRI Neurofeedback Based Stratification of Default Network Regulation Neuroimaging Data Repository

**DOI:** 10.1101/075275

**Authors:** Amalia R McDonald, Jordan Muraskin, Nicholas T Van Dam, Caroline Froehlich, Benjamin Puccio, John Pellman, Clemens CC Bauer, Alexis Akeyson, Melissa M Breland, Vince D Calhoun, Steven Carter, Tiffany P Chang, Chelsea Gessner, Alyssa Gianonne, Steven Giavasis, Jamie Glass, Steven Homan, Margaret King, Melissa Kramer, Drew Landis, Alexis Lieval, Jonathan Lisinski, Anna Mackay-Brandt, Brittny Miller, Laura Panek, Hayley Reed, Christine Santiago, Eszter Schoell, Richard Sinnig, Melissa Sital, Elise Taverna, Russell Tobe, Kristin Trautman, Betty Varghese, Lauren Walden, Runtang Wang, Abigail B Waters, Dylan Wood, F Xavier Castellanos, Bennett Leventhal, Stanley J Colcombe, Stephen LaConte, Michael P Milham, R Cameron Craddoc

## Abstract

This data descriptor describes a repository of openly shared data from an experiment to assess inter-individual differences in default mode network (DMN) activity. This repository includes cross-sectional functional magnetic resonance imaging (fMRI) data from the Multi Source Interference Task, to assess DMN deactivation, the Moral Dilemma Task, to assess DMN activation, a resting state fMRI scan, and a DMN neurofeedback paradigm, to assess DMN modulation, along with accompanying behavioral and cognitive measures. We report technical validation from n=125 participants of the final targeted sample of 180 participants. Each session includes acquisition of one whole-brain anatomical scan and whole-brain echo-planar imaging (EPI) scans, acquired during the aforementioned tasks and resting state. The data includes several self-report measures related to perseverative thinking, emotion regulation, and imaginative processes, along with a behavioral measure of rapid visual information processing.

Technical validation of the data confirms that the tasks deactivate and activate the DMN as expected. Group level analysis of the neurofeedback data indicates that the participants are able to modulate their DMN with considerable inter-subject variability. Preliminary analysis of behavioral responses and specifically self-reported sleep indicate that as many as 73 participants may need to be excluded from an analysis depending on the hypothesis being tested.

## 1 Introduction

The default mode network (DMN) consists of an aggregation of brain regions that are active during rest, as measured by BOLD signal, and are associated with spontaneous thought and emotion regulation [1, 2, 3]. The network is also commonly deactivated during cognitively demanding tasks [4]. Alterations to the DMN have been associated with a broad array of neuropsychiatric conditions [5, 6]. However, as the DMN is most commonly assessed during rest [7] or as a result of deactivation during a task [8], most studies fail to differentiate ability to modulate the DMN from the tendency to do so. Individuals likely vary both in their capacity to activate and/or deactivate the DMN and in spontaneous implementation of function related to the DMN. Just as the use of specific emotion regulation strategies (e.g., cognitive reappraisal) relate to psychopathology specifically in the tendency to use them vs. instructed use [9], the ability to modulate the DMN and tendency to do so may represent distinct and important domains of neural and psychological function [10]. Consistent with the National Institute of Mental Health’s Research Domain Criteria project [11], the ability and tendency to regulate the DMN may have both general and specific illness implications. For example, it could be that deficits in the ability to suppress DMN-related activity such as mind-wandering may be related to cognitive deficits across differing forms of mental illness [12], while an increased likelihood to engage in specific forms of mind-wandering (e.g., worry, rumination) may be more specific to anxiety and depression [13, 12, 14].

While psychological functions related to the DMN can be targeted, the DMN itself is somewhat more difficult to target, as the functions it instantiates are presumably multiply determined (i.e., invoking specific aspects of the DMN and/or other networks) and varied (i.e., different in nature and possibly kind) [15]. Recent advances in real-time fMRI (rt-fMRI) [16, 17, 18] have made it possible to provide participant-specific feedback about neural networks. These advances permit the addition of instructions to modulate given neural networks as well as the assessment of an individual’s ability to follow the instructions or modulate the specific network. In addition to collecting task-based and resting-state data, using rt-fMRI as neurofeedback may be critical to acquiring knowledge about tendencies and capability to regulate the DMN.

The Default Network Regulation Neuroimaging Repository contains data from a suite of fMRI experiments aimed at better understanding individual variation in DMN activity and modulation. Although it is a separate project, it has been harmonized with, and is distributed alongside, the Enhanced Nathan Kline Institute-Rockland Sample (NKI-RS) [19], which aims to capture deep and broad phenotyping of a large community-ascertained sample. In addition to the NKI-RS protocol, this project includes data collection from tasks that activate (Moral Dilemma task [8]) and deactivate the DMN (Multi-Source Interference Task [20]), a resting state scan, a novel real-time fMRI neurofeedback-based paradigm that specifically probes DMN modulation [16], and additional self-report measures.

In this data descriptor, we provide an overview of planned data collection, methods used, summaries of data collected and available to date, and validation analyses. New data will be released on a regular basis and will be available at the Collaborative Informatics and Neuroimaging Suite (COINS) Data Exchange (http://coins.mrn.org/dx) [21, 22], as well as the Neuroimaging Informatics Tools and Resources Clearinghouse (NITRC; http://www.nitrc.org/) and in the Amazon Web Services (https://aws.amazon.com/s3/).

## 2 Organization and access to the repository

Datasets for the present project can be accessed through the COINS Data Exchange (http://coins.mrn.org/dx) [21, 22], NITRC (http://fcon_1000.projects.nitrc.org/indi/enhanced/download.html) or through the Amazon Web Services S3 bucket (https://aws.amazon.com/s3/). Documentation on downloading the datasets can be found at http://fcon_1000.projects.nitrc.org/indi/enhanced/sharing.html. Data are available through both COINS and NITRC in the form of .tar files, containing all imaging and phenotypic data. The COINS data exchange (only accessible with DUA) offers a query builder tool, permitting the user to target and download files by specific search criteria (e.g., full completion of certain phenotypic measures and/or imaging sequences). Data are available through AWS as individual compressed NIfTI files that are arranged in the brain imaging data structure (BIDS) file structure [23].

All of the presented data are shared in accordance with the procedures of the NKI-RS study (http://fcon_1000.projects.nitrc.org/indi/enhanced/sharing.html). While a goal of the project is to maximize access to these data, privacy for the participants and their personal health information is paramount. De-identified neuroimaging data along with limited demographic information can be downloaded from the repository without restriction. To protect participant privacy, access to the high dimensional phenotypic and assessment data requires a data usage agreement (DUA). The DUA is relatively simple and requires a signature by an appropriate institutional representative. The DUA is available via Neuroimaging Informatics Tools and Resources Clearinghouse (Data Citation A1: http://fcon_1000.projects.nitrc.org/indi/enhanced/data/DUA.pdf).

### 2.1 Phenotypic data

Basic phenotypic data, which includes age, sex and handedness are available in the **participants.tsv** file at the root of the repository. Comprehensive phenotypic data are available as comma separated value (.csv) files from COINS after completing a minimal data usage agreement. Summary scores calculated from the RVIP task info area available through COINs and trial-by-trial response information is available via NITRC (Data Citation Y1: http://fcon_1000.projects.nitrc.org/indi/enhanced/RVIP-master.zip).

### 2.2 Imaging data

The imaging data is released in unprocessed form except that image headers have been wiped of protected health information and faces have been removed from the structural images. Data are available in NIfTI files arranged in the BIDS format [23]. Acquisition parameters are provided in JSON files that are named the same as their corresponding imaging data. Task traces (including stimulus onsets and durations) and responses, and physiological recordings, are also provided in JSON files along with the data. Additional details for all MRI data, as well as corresponding task information are available via NITRC (Data Citation X1: http://fcon_1000.projects.nitrc.org/indi/enhanced/mri_protocol.html#scans-acquired).

### 2.3 Quality assessment data

Quality metrics are available for download from the QAP repository (http://preprocessed-connectomes-project.org/quality-assessment-protocol). They are available as two (anatomical and functional) comma separated value files.

## 3 Contents of the Repository

The Neurofeedback (NFB) repository contains neuroimaging and assessment data that characterizes DMN function in a community ascertained sample of adults (21-45 years old) with a variety of psychiatric diagnoses. The data is collected in a separate 2.5-hour visit that occurred within **\*\*six\*\*** months of completing the Enhanced Nathan Kline Institute-Rockland Sample (NKI-RS) protocol [19]. The NKI-RS entails a 1 to 2-visit deep phenotyping protocol [19] and a Connectomes oriented neuroimaging assessment (data collection visit schedules are available online at: http://fcon_1000.projects.nitrc.org/indi/enhanced/sched.html). NKI-RS phenotyping includes a variety of cognitive and behavioral assessments, a blood draw, a basic fitness assessment, nighttime actigraphy, medical history questionnaire, and the administration of the Structured Clinical Interview for DSM-IV-TR Non-patient edition [24]. The NKI-RS imaging protocol includes structural MRI, diffusion MRI, several resting state fMRI, and perfusion fMRI scans. Although the NKI-RS data is being openly shared (http://fcon_1000.projects.nitrc.org/indi/enhanced/) it is not a part of the NFB repository and is not described in further detail herein.

### 3.1 Participants

The NFB repository will ultimately contain data from a total of 180 residents of Rockland, Westchester, or Orange Counties, New York, or Bergen County, New Jersey aged 21 to 45, (50% male at each age year). Based on census data from 2013, Rockland County, New York has the following demographics [25]: Median age 36.0 years, population is 50.5% female, 80.4% White, 11.4% Black/African American, 6.3% Asian, 0.5% Native American/Pacific Islander, and 1.4% “Other”. With regards to ethnicity, 13.8% of the population endorses being Hispanic/Latino.

Minimally restrictive psychiatric exclusion criteria, which only screen out severe illness, were employed to include individuals with a range of clinical and sub-clinical psychiatric symptoms. Medical exclusions include: chronic medical illness, history of neoplasia requiring intrathecal chemotherapy or focal cranial irradiation, premature birth (prior to 32 weeks estimated gestational age or birth weight < 1500g, when available), history of neonatal intensive care unit treatment exceeding 48 hours, history of leukomalacia or static encephalopathy, or other serious neurological (specific or focal) or metabolic disorders including epilepsy (except for resolved febrile seizures), history of traumatic brain injury, stroke, aneurysm, HIV, carotid artery stenosis, encephalitis, dementia or mild cognitive impairment, Huntington’s Disease, Parkinson’s, hospitalization within the past month, contraindication for MRI scanning (metal implants, pacemakers, claustrophobia, metal foreign bodies or pregnancy) or inability to ambulate independently. Severe psychiatric illness can compromise the ability of an individual to comply with instructions, tolerate the MRI environment and participate in the extensive phenotyping protocol. Accordingly, participants with severe psychiatric illness were excluded, as determined by Global Assessment of Function (GAF; DSM-IV TR [24]) < 50, history of chronic or acute substance dependence disorder, history of diagnosis with schizophrenia, history of psychiatric hospitalization, or suicide attempts requiring medical intervention.

The technical validation reported in this paper includes 125 participants with the following demographics: median age 30, mean age, 31, std. dev. 6.6, 77 females, 59.2% White, 28.8% Black/African American, 0.56% Asian, 0% Native American/Pacific Islander, and 0.56% “Other”, and 16.8% Hispanic/Latino. A summary of participant diagnoses and current medications are listed in tables 1 and 2.

**Table 1.** Medications used by participants on a daily basis. Medications were grouped by class and the number of participants who reported taking them is provided (Pts.). Full medication information regarding drug name, dosage, primary indication, and duration of time taken at a participant level can be accessed through COINs after completing a DUA.

| Medication Class | Pts. |
| --- | --- |
| Vitamins/Supplements | 34 |
| Obstetrics/Gynecology | 12 |
| Psychiatric | 8 |
| Gastrointestinal | 6 |
| Endocrine | 5 |
| Analgesics | 4 |
| Cardiovascular | 3 |
| Rheumatologic | 2 |
| Asthma/Pulmonary | 2 |
| Allergy/Cold/ENT | 1 |
| Immunology | 1 |
| Dermatologic | 1 |
| Antihistamine | 1 |

**Table 2.** Summary diagnostic information for the included 125 participants as determined by the SCID-IV or consensus diagnosis by the study psychiatrist. Diagnoses are accompanied with the SCID diagnostic code and the number of participants diagnosed (Pts.). Full diagnostic information

| Diagnosis (SCID-IV code) | Pts. |
| --- | --- |
| No Diagnosis or Condition on Axis I (V71.09) | 59 |
| Alcohol Abuse Past (305.00) | 21 |
| Major Depressive Disorder, Single Episode, In Full Remission (296.26) | 12 |
| Cannabis Abuse Current (305.20) | 12 |
| Cannabis Dependence Past (304.30) | 11 |
| Major Depressive Disorder, Recurrent, In Full Remission Past (296.36) | 8 |
| Major Depressive Disorder, Single Episode, Unspecified Past (296.20) | 5 |
| Posttraumatic Stress Disorder Current (309.81) | 5 |
| Alcohol Dependence Past (303.90) | 5 |
| Specific Phobia Past (300.29) | 5 |
| Generalized Anxiety Disorder Current (300.02) | 4 |
| Attention-Deficit/Hyperactivity Disorder, Inattentive Type Current (314.00) | 4 |
| Attention-Deficit/Hyperactivity Disorder NOS Current (314.9) | 3 |
| Attention-Deficit/Hyperactivity Disorder, Hyperactive-Impulsive Type Current (314.01) | 3 |
| Cocaine Abuse Past (305.60) | 2 |
| Anorexia Nervosa Past (307.1) | 2 |
| Anxiety Disorder NOS Current (300.00) | 2 |
| Panic Disorder Without Agoraphobia Past (300.01) | 2 |
| Social Phobia Current (300.23) | 2 |
| Agoraphobia Without History of Panic Disorder Current (300.22) | 2 |
| Panic Disorder With Agoraphobia Past (300.21) | 2 |
| Hallucinogen Abuse Past (305.30) | 2 |
| Cocaine Dependence Past (304.20) | 2 |
| Trichotillomania (312.39) | 1 |
| Bulimia Nervosa Current (307.51) | 1 |
| Major Depressive Disorder, Recurrent, In Partial Remission Past (296.35) | 1 |
| Major Depressive Disorder, Recurrent, Moderate Current (296.32) | 1 |
| Bereavement (V62.82) | 1 |
| Hallucinogen Dependence Past (304.50) | 1 |
| Obsessive-Compulsive Disorder Current (300.3) | 1 |
| Body Dysmorphic Disorder Current (300.7) | 1 |
| Eating Disorder NOS Past (307.50) | 1 |
| Phencyclidine Abuse Past (305.90) | 1 |
| Delusional Disorder Mixed Type (297.1) | 1 |
| Amphetamine Dependence Past (304.40) | 1 |
| Opioid Abuse Past (305.50) | 1 |
| Sedative, Hypnotic, or Anxiolytic Dependence Past (304.10) | 1 |

### 3.2 Phenotypic Data

In addition to the NKI-RS protocol, participants completed a variety of assessments that probe cognitive, emotional, and behavioral domains that have been previously implicated with DMN function [2, 13, 12, 3]. These included the Affect Intensity Measure (AIM) [26]–to measure the strength or weakness with which one experiences both positive and negative emotions, Emotion Regulation Questionnaire (ERQ; [27])–to assess individual differences in the habitual use of two emotion regulation strategies: cognitive reappraisal and expressive suppression, Penn State Worry Questionnaire (PSWQ; [28])–to measure worry, Perseverative Thinking Questionnaire (PTQ; [29])–to measure the broad idea of repetitive negative thought, Positive and Negative Affect Schedule - short version (PANAS-S; [30])–to measure degrees of positive or negative affect, Ruminative Responses Scale (RRS; [31])–to assess rumination that is related to, but not confounded by depression, and the Short Imaginal Process Inventory (SIPI; [32])–to measure aspects of daydreaming style and content, mental style, and general inner experience. Assessments were completed using web-based forms implemented in COINs [21]. Sample mean, standard deviation, and range of the above measures for the first 125 participants are provided in Table 3.

**Table 3.**
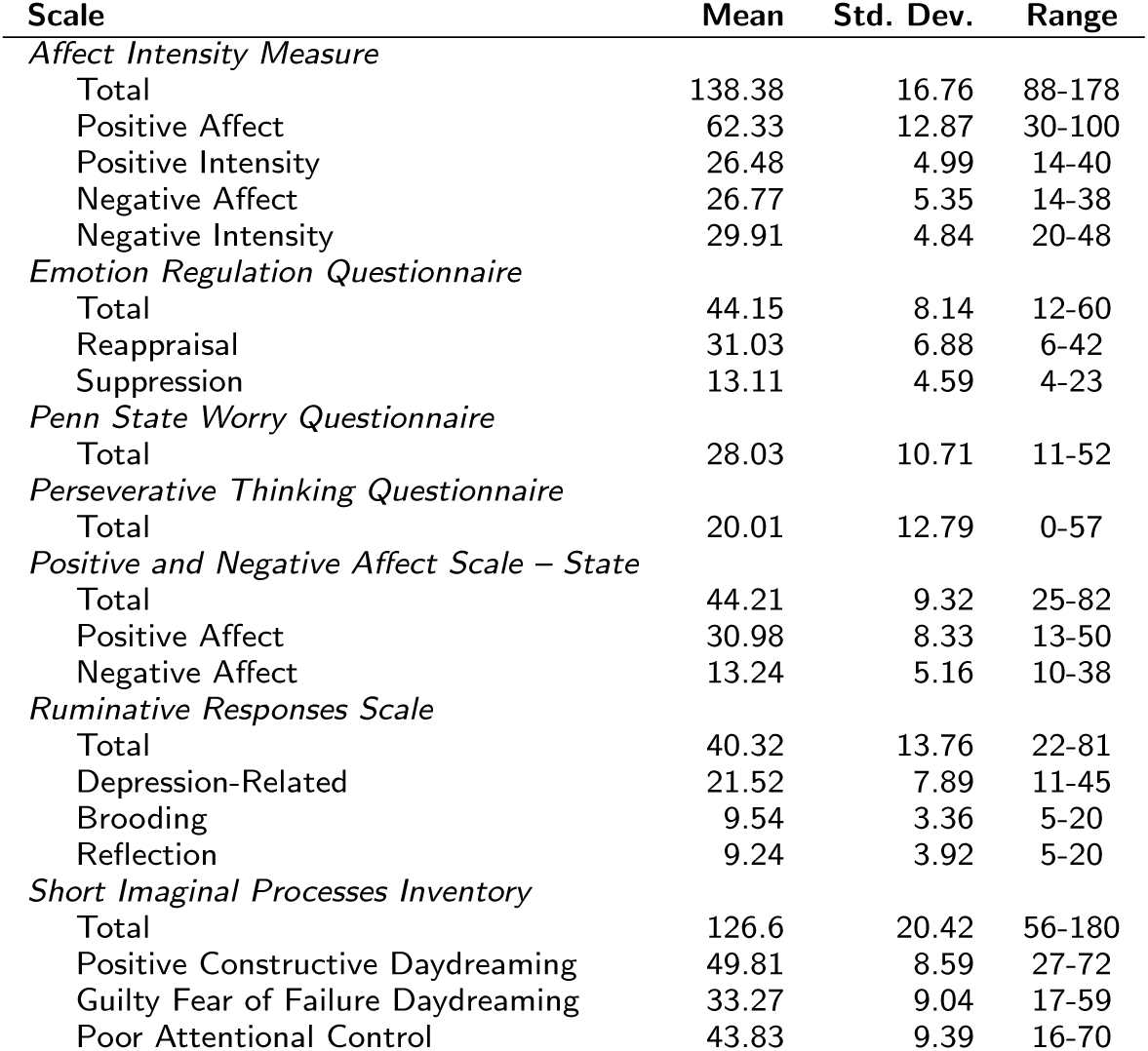
Statistics for self-report measures from the neurofeedback specific visit related to perseverative thinking, emotion regulation, and imaginative processes. Mean, Standard Deviation, and Range are reported for the total AIM, ERQ, PSWQ, PTQ, PANAS, RRS, and SIPI scores, as well as each measure’s sub-scales.

Participants also completed a Rapid Visual Information Processing (RVIP) task to assess sustained attention and working memory. Response times and detection accuracy from this task have been previously correlated with DMN function [33]. The RVIP is administered using custom software, implemented in PsychoPy [34], which conforms to the literature describing its original implementation in the Cambridge Neuropsychological Test Automated Battery (CANTAB [35]). A pseudorandom stream of digits (0-9) is presented to the participants in white, centered on a black background, surrounded by a white box. Participants are instructed to press the space bar whenever they observe the sequences 2-4-6, 3-5-7, or 4-6-8. Digits are presented one after another at a rate of 100 digits per minute and the number of stimuli that occurred between targets varied between 8 and 30. Responses that occurred within 1.5 seconds of the last digit of a target sequence being presented were considered “hits”. Stimuli presentation continued until a total of 32 target sequences were encountered, which required on average 4 minutes and 20 seconds. Before performing the task, participants completed a practice version that indicated when a target sequence had occurred and provided feedback (“hit” or “false alarm”) whenever the participant pressed the space bar. Participants were allowed to repeat the practice until they felt that they were comfortable with the instructions. Summary statistics calculated from the RVIP included: mean reaction time, total targets, hits, misses, false alarms, hit rate (*H*), false alarm rate (*F*), and *A’* (Eqn. 1), *A’* is an alternative to the more common d’ in signal detection theory [36]). The task can be downloaded from the OpenCogLab Repository (http://opencoglabrepository.github.io/experiment_rvip.html).

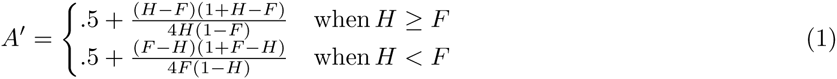

### 3.3 MRI Acquisition

Data was acquired on a 3 T Siemens Magnetom TIM Trio scanner (Siemens Medical Solutions USA: Malvern PA, USA) using a 12-channel head coil. Anatomic images were acquired at 1 **×** 1 **×** 1 mm^3^ resolution with a 3D T1-weighted magnetization-prepared rapid acquisition gradient-echo (MPRAGE) sequence [37] in 192 sagittal partitions each with a 256 **×** 256 field of view (FOV), 2600 ms repetition time (TR), ms echo time (TE), 900 ms inversion time (TI), 8° flip angle (FA), and generalized auto-calibrating partially parallel acquisition (GRAPPA) [38] acceleration factor of 2 with 32 reference lines. The sMRI data were acquired immediately after a fast localizer scan and preceded the collection of the functional data.

A gradient echo field map sequence was acquired with the following parameters: TR 500 ms, TE1/TE2 2.72 ms/5.18 ms, FA 55°, 64 **×** 64 matrix, with a 220 mm FOV, 30 3.6 mm thick interleaved, oblique slices, and in plane resolution of 3.4 **×** 3.4 mm^2^. All functional data were collected with a blood oxygenation level dependent (BOLD) contrast weighted gradient-recalled echo-planar-imaging sequence (EPI) that was modified to export images, as they were acquired, to AFNI over a network interface [39, 40]. FMRI acquisition consisted of 30 3.6mm thick interleaved, oblique slices with a %10 slice gap, TR 2000 ms, TE 30 ms, FA 90°, 64 **×** 64 matrix, with 220 mm FOV, and in plane resolution of 3.4 **×** 3.4 mm^2^. Functional MRI scanning included a three volume “mask” scan and a six-minute resting state scan followed by three task scans (described later) whose order was counterbalanced across subjects.

During all scanning, galvanic skin response (GSR), pulse and respiration waveforms were measured using MRI compatible non-invasive physiological monitoring equipment (Biopac Systems, Inc.). Rate and depth of breathing were measured with a pneumatic abdominal circumference belt. Pulse was monitored with a standard infrared pulse oximeter placed on the tip of the index finger of the non-dominant hand. Skin conductance was measured with disposable passive electrodes that were non-magnetic and non-metallic, and collected on the hand. The physiological recordings were synchronized with the imaging data using a timing signal output from the scanner. Visual stimuli were presented to the participants on a projection screen that they could see through a mirror affixed to the head coil. Audio stimuli were presented through headphones using an Avotec Silent-Scan^®^ pneumatic system (Avotec, Inc.: Stuart FL, USA).

### 3.4 MRI acquisition order and online Processing

The real time fMRI neurofeedback experiment utilizes a classifier based approach for extracting DMN activity levels from fMRI data TR-by-TR, similar to [16]. The classifier is trained from resting state fMRI data using a time course of DMN activity extracted from the data using spatial regression to a publicly available template derived from a meta-analysis of task and resting state datasets [41, 42]. Several stages of online processing are necessary to perform this classifier training, as well as, denoising of fMRI data in real-time. These stages include calculating transforms required to convert the DMN template from MNI space to subject space, creating white matter (WM) and cerebrospinal fluid (CSF) masks for extracting nuisance signals, and training a support vector regression (SVR) model for extracting DMN activity. The MRI session was optimized to collect the data required for these various processing steps, and to perform the processing, while minimizing delays in the experiment.

After acquiring a localizer, the scanning protocol began with the acquisition of a T1 weighted anatomical image used for calculating transforms to MNI space and white matter and CSF masks. Once the image was acquired it was transferred to a DICOM server on the real-time analysis computer (RTAC), which triggered initialization of online processing. The processing script started AFNI in real-time mode, configured it for fMRI acquisition, and began structural image processing. Structural processing included reorienting the structural image to RPI voxel order, skull-stripping using AFNI’s 3dSkullStrip [43], resampling the image to isotropic 2mm voxels (to reduce computational cost of subsequent operations), segmentation into grey matter (GM), white matter (WM), and cerebrospinal fluid (CSF) using FSL’s FAST [44], and normalization into MNI space using FSL’s FLIRT [45, 46]. WM and CSF probability maps were binarized using a 90% threshold. The CSF mask was constrained to the lateral ventricles using an ROI from the AAL atlas to avoid overlap with GM.

In parallel with the structural processing a FieldMap was collected but not used in the online processing. Subsequently, a three volume “mask” EPI scan was acquired and transferred to the RTAC. The three images were averaged, reoriented to RPI, and used to create a mask to differentiate brain signal from background (using AFNI’s 3dAutomask [43]). The mean image was coregistered to the anatomical image using FSL’s boundary based registration (BBR) [47]. The resulting linear transform was inverted and applied to the WM and CSF masks to bring them into alignment with the fMRI data. Additionally, the transform was combined with the inverted anatomical-to-MNI transform and applied to the canonical map of the DMN (from [41]).

Next, a 6-minute resting state scan (182 volumes) was collected and used as training data to create the support vector regression (SVR) model. This procedure involved motion correction followed by a nuisance regression procedure to orthogonalize the data to six head motion parameters, mean WM, mean CSF, and global signal [48, 49, 50]. A SVR model of the DMN was trained using a modified dual regression procedure in which a spatial regression to the unthresholded DMN template was performed to extract a time course of DMN activity. The Z-transformed DMN time course was then used as labels (independent variable), with the preprocessed resting state data as features (dependent variables), for SVR training (C=1.0, e = 0.01) using AFNI’s 3dsvm tool [51]. The result was a DMN map tailored to the individual participant based on preexisting expectations about DMN anatomy and function. After SVR training was completed (generally took less than 2 minutes), the MSIT (198 volumes), Moral Dilemma (144 volumes), and Neurofeedback test (412 volumes) scans were run. The order of the task based functional scans was counterbalanced and stratified for age and sex across participants.

### 3.5 Functional MRI Tasks

#### 3.5.1 Resting state scan

Participants were instructed to keep their eyes open during the scan and fixate on a white plus (+) sign centered on a black background. They were asked to let their mind wander freely and if they noticed themselves focusing on any particular train of thought, to let their mind wander away from it.

#### 3.5.2 Moral Dilemma Task

The Moral Dilemma (MD) task involved a participant making a decision about what he or she would do in a variety of morally ambiguous scenarios presented as a series of vignettes (see Fig. 1A for an example) [8]. As a control, the participant was asked to recall a detail from a dilemma-free vignette (see Fig. 1B for an example). Prior work using these vignettes has shown strong activation of DMN regions during moral dilemma scenarios relative to control scenarios [8]. The MD task provided a basis to directly examine task-induced activation of the DMN.

**Figure 1.**
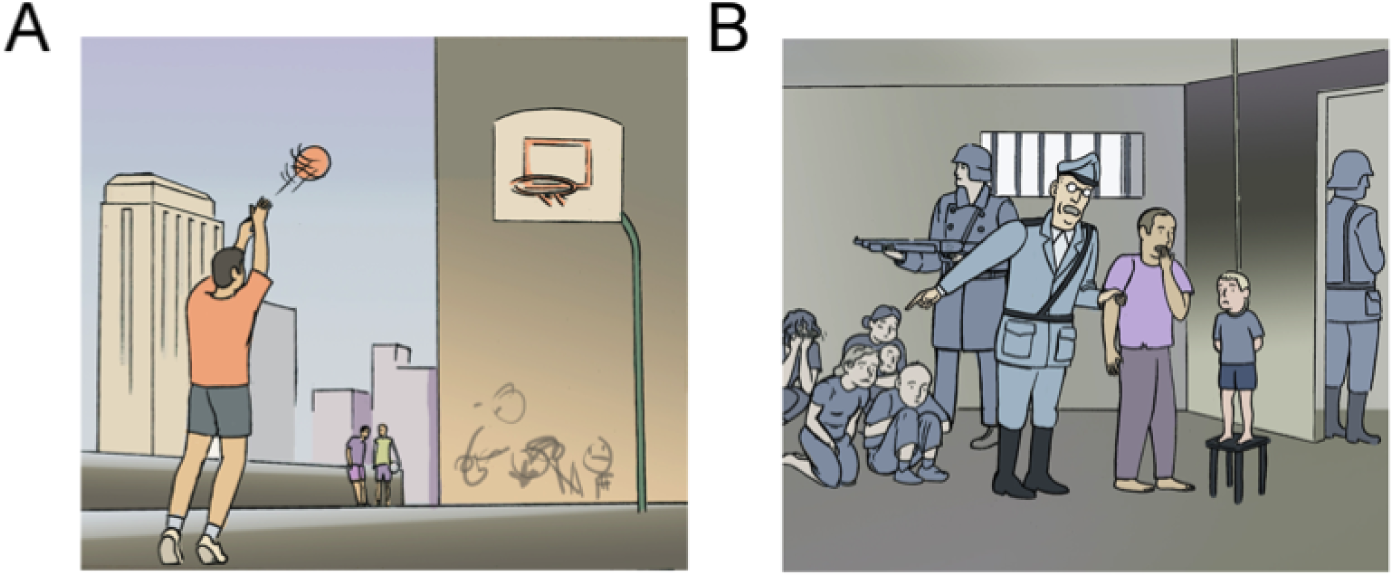
Examples of vignettes from the moral dilemma task. A) A control vignette: “Mr. Jones is practicing his three-point throw on the basketball court behind his house. He hasn’t managed to score a basket during the whole morning, despite all the practice. He concentrates hard and throws the ball one more time. This time his aim is more accurate, the ball curves through the air and falls cleanly into the basket. Mr. Jones has managed to score a basket for the first time.” Question during task: Will he score? B) A dilemma vignette. “Mr. Jones and his only son are held in a concentration camp. His son tries to escape but he is caught. The guard watching them tells Mr. Jones that his son is going to be hanged and that it will be him (Mr. Jones) who has to push the chair. If he does not do it, not only will his son die but also five more people held in the concentration camp.” Question during task: Would you push the chair? Vignettes are copied from [8] Support Information Appendix.

Just prior to the scan, participants listened to a recording of the vignette while viewing a corresponding image (see Fig. 1A and 1B for examples) and were asked to decide how they would react in the scenario. The fMRI tasks consisted of 24 moral dilemma questions and 24 control questions presented in eight 30-second blocks, each consisting of six questions, that alternated between moral dilemma and control conditions. Participants viewed an image and heard a question corresponding to each vignette. Each image was displayed for 5 seconds and the audio began one second after image onset. Participants responded to the proposed question by pressing one of two buttons on a response box (index finger button for “yes”, middle finger button for “no”). The task began and ended with 20 second fixation blocks during which participants passively viewed a plus (+) sign centered on a grey background. The task was implemented in PsychoPy [34] using images provided by the authors of the original work [8] and can be downloaded from the OpenCogLab Repository (http://opencoglabrepository.github.io/experiment_moraldillema.html).

#### 3.5.3 Multi-Source Interference Task

The MSIT was developed as an all-purpose task to provide robust single-participant level activation of cognitive control and attentional regions in the brain [20]. Early work with the MSIT suggests robust activation of regions associated with top-down control – regions that are often active when the DMN is inactive [20]. The MSIT provided a basis for directly examining task-induced deactivation of the DMN.

During the task, participants were presented with a series of stimuli consisting of three digits, two of which were identical and one that differed from the other two (see Fig. 2 for examples). Participants were instructed to indicate the value and not the position of the digit that differed from the other two (e.g., 1 in 100, 1 in 221). During control trials, distractor digits were 0s and the targets were presented in the same location as they appear on the response box (e.g., 1 in 100 was the first button on the button box and the first number in the sequence). During interference trials, distractors were other digits and target digits were incongruent with the corresponding location on the button box (e.g., 221 – 1 was the first button on the button box but was the third number in the sequence).

**Figure 2.**
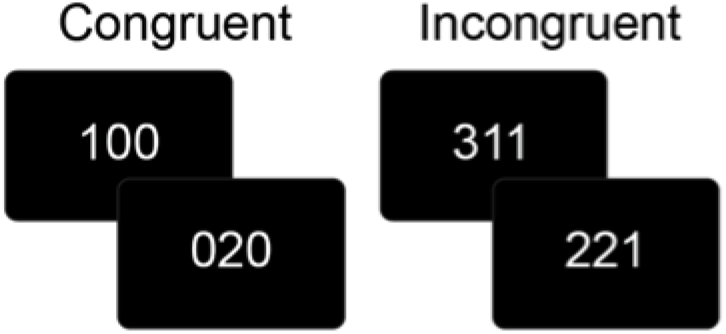
Examples of MSIT task stimuli. For congruent trials, the paired digits are zero and the position of the target digit corresponds to its location. For incongruent trials, the distractor digits are non-zero and the target digits location is not the same as its value.

The task was presented as a block design with eight 42-second blocks that alternated between conditions, starting with a control block. Each block contained 24 randomly generated stimuli with an inter-stimulus interval of 1.75 seconds. The task began and ended with a 30 second fixation period during which they passively viewed a white plus (+) sign centered on a black background. The MSIT was implemented in PsychoPy (Peirce, 2008) and can be downloaded from the OpenCogLab Repository (http://opencoglabrepository.github.io/experiment_msit.html).

#### 3.5.4 Neurofeedback Task

In the real-time Neurofeedback (rt-NFB) task, fMRI data were processed during collection, allowing the experimenter to provide visual feedback of brain activity over the course of the experiment [39, 17]. The rt-NFB task was developed to examine each individual participant’s ability to either increase or decrease DMN activity in response to instructions, accompanied by real-time feedback of activity from their own DMN [16].

During the rt-NFB task, participants were shown an analog meter with ‘Wander’ at one end and ‘Focus’ at the other, along with an indicator of their current performance (see Fig. 3). The fixation point was a white square positioned equally between the two poles. Participants were instructed at the beginning of each block to attempt to either focus their attention (Focus) or to let their mind wander (Wander). The task began with a 30 second control condition and then proceeded with alternating blocks with a specific sequence of durations – 30, 90, 60, 30, 90, 60, 60, 90, 30, 60, 90, and 30 seconds. At the end of each block, the participant was asked to press a button within a two second window. Starting condition (‘Focus’ vs. ‘Wander’) was counterbalanced, as was the location of each descriptor (‘Focus’, ‘Wander’; Right vs. Left) on the analog meter was counterbalanced across participants.

**Figure 3.**
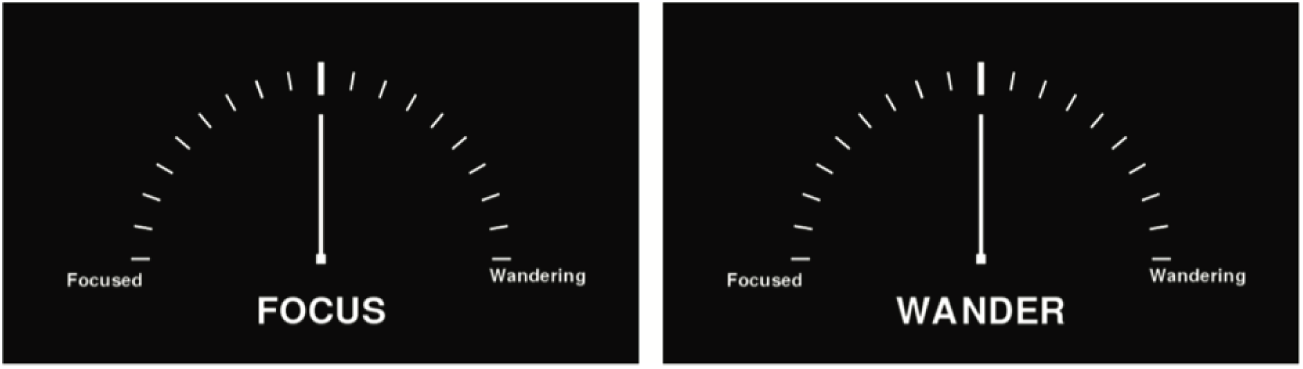
Stimuli for “Focus” and “Wander” conditions of the neurofeedback task. The needle moves to the left or the right based on DMN activity.

Each fMRI volume acquired during the task was transmitted from the scanner to the RTAC shortly after it was reconstructed and passed to AFNI’s real-time plugin [39] for online processing. The volume was realigned to the previously collected mean volume to correct for motion and to bring it into alignment with the tissue masks and DMN SVR model. Mean WM intensity, mean CSF intensity, and global mean intensity were extracted from the volume using masks calculated from the earlier online segmentation of the anatomical image. A general linear model was calculated at each voxel, using all of the data that were acquired up to the current time point, with WM, CSF, global, and motion parameters included as regressors of no interest. The recently acquired volume was extracted from the residuals of the nuisance variance regression, spatially smoothed (6-mm FWHM), and then a measure of DMN activity was decoded from the volume using the SVR model trained from the resting state data. The resulting measure of DMN activity was translated to an angle, which was added to the current position of the needle on the analog meter, moving it in the direction of focus or wander based on DMN activation or deactivation, respectively. This moving average procedure was used to smooth the motion of the needle. The position of the needle was reset to the center at each change between conditions. The neurofeedback stimulus was implemented in Vision Egg [52] and can be downloaded from the OpenCogLab Repository (http://opencoglabrepository.github.io/experiment_RTfMRIneurofeedback.html).

### 3.6 Quality Measures

Metrics of the spatial quality of the structural and functional MRI data and temporal quality of the fMRI data were calculated using the Preprocessed Connectomes Project Quality Assessment Protocol (QAP; http://preprocessed-connectomes-project.org/quality-assessment-protocol, [53]). For the structural data, spatial measures include:

- Signal-to-Noise Ratio (SNR): mean grey matter intensity divided by standard deviation of out-of-brain (OOB) voxels, larger values are better [54].
- Contrast-to-Noise Ratio (CNR): the difference between mean gray matter intensity and mean white matter intensity divided by standard deviation of OOB voxels, larger values are better [54].
- Foreground-to-Background Energy Ratio (FBER): variance of in-brain (IB) voxels divided by variance of OOB voxels, larger values are better.
- Percent artifact voxels (QI1): number of OOB voxels that are in structured noise (i.e. artifacts) divided by the total number of OOB voxels, smaller values are better [55].
- Spatial smoothness (FWHM): smoothness of voxels expressed as full-width half maximum of the spatial distribution of voxel intensities in units of voxels, smaller values are better [56].
- Entropy focus criterion (EFC): the Shannon entropy of voxel intensities normed by the maximum possible entropy for an image of the same size, is a measure of ghosting and blurring induced by head motion, lower values are better [57]
- Summary measures: mean, standard deviation, and size of different image compartments, including, foreground, background, WM, GM, and CSF.

Spatial measures of fMRI data include EFC, FBER, FWHM, and well as:

- Ghost-to-signal Ratio (GSR): mean of the voxel intensities in parts of the fMRI image (determined by the phase encoding direction) that are susceptible to ghosting, smaller values are better [58].
- Summary measures: mean, standard deviation, and size of foreground and background.

Temporal measures of fMRI data include:

- Standardized DVARS (DVARS): the mean intensity change between every pair of consecutive fMRI volumes, standardized to make it comparable between scanning protocols, smaller values are better [59].
- Outliers: the mean number of outlier voxels found in each volume using AFNI’s 3dToutcount command, smaller values are better [43].
- Global correlation (gcorr): the average correlation between every pair of voxel time series in the brain, sensitive to physiological noise, head motion, as well as imaging technical artifacts, such as, signal bleeding in multi-band acquisitions, values that are closer to zero are better [60].
- Mean root mean square displacement (mean RMSD): the average distance between consecutive fMRI volumes [61]. This has shown to be a more accurate representation of head motion than mean frame-wise displacement (meanFD) proposed by Power et. al. [62] [63].

## 4 Technical Validation

A variety of initial analyses have been performed on the first 125 participants to be released in the NFB repository to establish the quality of these data and the successful implementation of the tasks. A series of quality assessment measures were calculated from the raw imaging data and compared with data available through other data sharing repositories. Results of the behavioral tasks were evaluated to ensure consistency with the existing literature. Preliminary analyses of the various fMRI tasks were performed to verify activation and deactivation of the DMN as predicted.

### 4.1 Behavioral Assessment

Participant responses for the RVIP, MSIT, moral dilemma task and neurofeedback task were analyzed to evaluate whether the participants complied with task instructions, and whether their responses are consistent with previous literature.

#### RVIP

The RVIP python script calculated hit rate, false alarm rate, *A’*, and mean reaction time during task performance. Responses that occur within 1.5 seconds of the last digit of a target sequence being displayed were considered hits, multiple responses within 1.5 seconds are considered a hit followed by multiple false alarms, and responses that occur outside of the 1.5-second window are considered false alarms. The number of hits and false alarms are converted to rates by dividing by the total number of targets. Since the number of false alarms are not bounded, the false alarm rate can be higher than 100%, resulting in *A’* values greater than 1 (see Eqn. 1). In post-hoc analysis, false alarm raters greater than 1 were replaced with 1 and *A’* values greater than 1 were replaced with 0.

#### MSIT

Reaction times and correctness of response were calculated for each trial by the MSIT python script. The response window begins at stimulus presentation and ends just prior to the presentation of the next stimulus. The last key pressed after stimulus presentation is considered in these calculations. A summary script distributed with the task was used to summarize response times and accuracy rates for congruent and incongruent trials. This script assumes that the task begins with a block of congruent stimuli.

#### Moral Dilemma

Reaction times and correctness of responses for control trials were calculated for each trial from the task log files using a script distributed with the task. The response window starts at the beginning of the auditory question prompt and the last key received in this window is used to calculate response times. The length of the auditory question was subtracted from reaction times to control for question length across stimuli. This may result in negative reaction times. These values were then summarized into average reaction time for control and dilemma stimuli. Response accuracy was calculated for the control trials, there is no definitively correct response to the dilemmas.

#### Self-report sleep

After completing the scans, participants were asked to list the scans that they fell asleep during, if any. The results are coded in the COINS database and were analyzed to determine which of the participants fell asleep during the training or testing scans.

#### Neurofeedback

Participants are asked to press a button at transitions between “focus” and “wander” blocks. The hit rate for these catch trials were calculated from the task log files using a script distributed with the task.

### 4.2 FMRI Analysis

All data were preprocessed using a development version of the Configurable Pipeline for the Analysis of Connectomes (C-PAC version 0.4.0, http://fcp-indi.github.io). C-PAC is an open-source, configurable pipeline for the automated preprocessing and analysis of fMRI data [64]. C-PAC is a software implemented in Python that integrates tools from AFNI [43], FSL [65], and ANTS [66] with custom tools, using the Nipype [67] pipelining library, to achieve high-throughput processing on high performance computing systems.

Anatomical processing began with skull stripping using the BEaST toolset [68] with a study-specific library and careful manual correction of the results. The masks and BEaST library generated through this effort are shared through the Preprocessed Connectomes Project NFB Skullstripped repository (http://preprocessed-connectomes-project.org/NFB_skullstripped/) [69]. Skull-stripped images were resampled to RPI orientation and then a non-linear transform between images and a 2mm MNI brain-only template (FSL, [65]) was calculated using ANTs [66]. The skullstripped images were additionally segmented into WM, GM, and CSF using FSL’s FAST tool [44]. A WM mask was calculated by applying a 0.96 threshold to the resulting WM probability map and multiplying the result by a WM prior map (avg152T1_white_bin.nii.gz - distributed with FSL) that was transformed into individual space using the inverse of the linear transforms previously calculated during the ANTs procedure. A CSF mask was calculated by applying a 0.96 threshold to the resulting CSF probability map and multiplying the result by a ventricle map derived from the Harvard-Oxford atlas distributed with FSL [70]. The thresholds were chosen and the priors were used to avoid overlap with grey matter.

Functional preprocessing began with resampling the data to RPI orientation, and slice timing correction. Next, motion correction was performed using a two-stage approach in which the images were first coregistered to the mean fMRI and then a new mean was calculated and used as the target for a second coregistration (AFNI 3dvolreg [71]). A 7 degree of freedom linear transform between the mean fMRI and the structural image using FSL’s implementation of boundary-based registration [47]. Nuisance variable regression (NVR) was performed on motion corrected data using a 2nd order polynomial, a 24-regressor model of motion [48], 5 nuisance signals, identified via principal components analysis of signals obtained from white matter (CompCor, [72]), and mean CSF signal. WM and CSF signals were extracted using the previously described masks after transforming the fMRI data to match them in 2mm space using the inverse of the linear fMRI-sMRI transform. NVR residuals were written into MNI space at 3mm resolution and subsequently smoothed using a 6mm FWHM kernel.

Individual level analyses of the MSIT and MD tasks were performed in FSL using a general linear model. The expected hemodynamic response of each task condition was derived from a boxcar model, specified from stimulus onset and duration times, convolved with a canonical hemodynamic response function. Multiple regressions were performed at each voxel with fMRI activity as the independent variable and task regressors as the dependent variables. Regression coefficients at each voxel were contrasted to derive a statistic for the difference in activation between task conditions (incongruent > congruent for MSIT, dilemma > control for MD). The resulting individual level task maps were entered into group-level one-sample t-tests, whose significance were assessed using multiple comparison correction via a permutation test (10,000 permutations) implemented by FSL’s randomise *(p <* 0.001 FWE - Threshold Free Cluster Enhancement (TFCE) [73]). Participants with missing behavioral data, or whose behavioral responses were outliers (> 1.5 interquartile range), were excluded from group level analysis.

Maps of DMN functional connectivity were extracted from the resting state and neurofeedback scans using a dual regression procedure [74]. Time courses from 10 commonly occurring intrinsic connectivity networks (ICNs) were extracted by spatially regressing each dataset onto templates derived from a meta-analysis of task and resting state datasets [41]. The resulting time courses were entered into voxel-wise multiple regressions to derive connectivity map for each of the 10 ICNs. The ICN corresponding to the DMN was subsequently subtracted and entered into group-level one-sample t-tests, whose significance were assessed using multiple comparison correction via a permutation test (10,000 permutations) implemented by FSL’s randomise (p < 0.001 FWE - Threshold Free Cluster Enhancement (TFCE) [73]). To evaluate performance on the neurofeedback task, DMN time course for each participant were correlated with an ideal time course (obtained by using instruction onset and duration information – i.e., focus or wander). Participants who reported sleeping during the resting-state scan were excluded from its analysis and analysis of the feedback scan was performed with and without participants who reported sleep.

### 4.3 Validation Results

#### 4.3.1 Behavioral Assessment

The behavioral results for the Rapid Visual Information Processing (RVIP), Multi-Source Interference Task (MSIT), Moral Dilemma Task (MD), and Neurofeedback Task (NFB) are illustrated in Fig. 4. Reaction times, hit rates, and A’ calculated from the RVIP task were consistent with, although slightly worse than, those previously published in young adults (n=20, age 25.30 +/- 5.09 years, RVIP reaction time 425.6 +/- 43.0 ms, A’ = 0.936 +/- 0.05, [75]). The differences in performance may have been due to the inclusion of participants with psychiatric diagnoses in the NFB population. The RVIP had the highest number of outliers of the tasks with 16 participants having false alarm rates greater than 1.5 times the inter-quartile range, 11 of which also had outlier A’ values (see Fig. 5). Similarly, Figure 4B illustrates that the performance on the MSIT was consistent with, although a little worse than, values published in a previous validation study (n = 8, 4 females, age = 30.4 +/- 5.6 years, congruent trial reaction time = 479 +/- 92 ms, incongruent trial reaction time = 787 +/- 129 ms [76]). MSIT data was only available in 107 of the participants due to a faulty button box, of which a total of 13 had responses that were outliers on at least one of the statistics derived from the task (see Fig. 5). For MD, the difference in reaction times distributions between control and dilemma trials support the hypothesis that the dilemma trials require more processing, as is expected [8] (see Fig. 4C). MD results are available for 121 participants due to button box problems and 12 of the remaining have accuracy on the control trials that are outliers (see Fig. 5). Only 58.5% of the participants reported remaining awake during both the resting state and neurofeedback scans, these subjective measures were validated by catch trial performance (see Fig. 4D).

**Figure 4.**
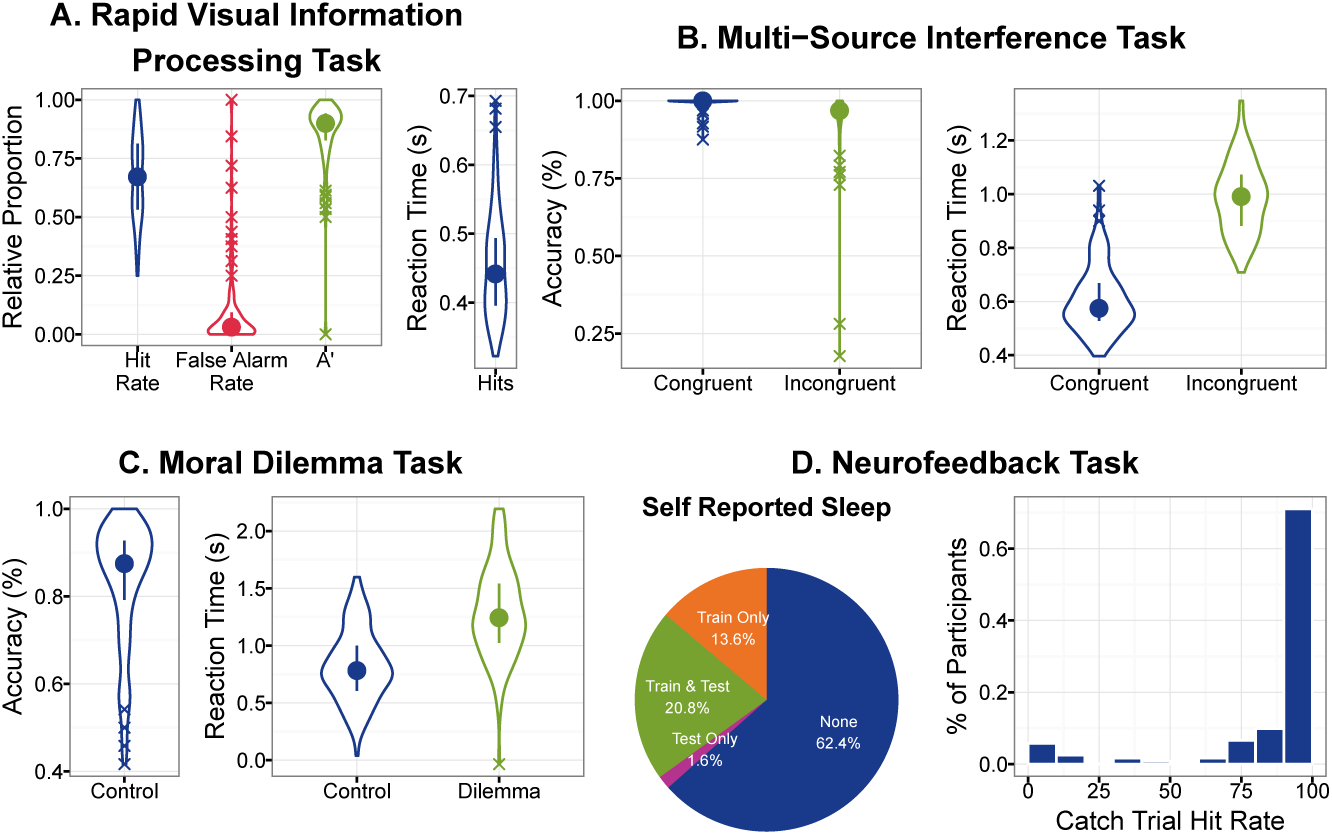
Visualizations of behavioral results for all tasks. Panels A-C show data as violin plots (filled circle represents the mean, line range represents the confidence interval, ‘X’ data points indicate outliers (1.5x Interquartile range)). Panel A shows, A’, False Alarm Rate, Hit Rate, and mean reaction time for the RVIP. Panel B shows accuracy (left) and reaction time (right) for congruent relative to incongruent trials of the MSIT task. Panel C shows accuracy for the control trials, as well as reaction time for both the dilemma and control trials of the Moral Dilemma Task. Panel D shows self-reported sleep (none, sleep during train only, sleep during test only, and sleep during train and test sessions) as a percentage of the total sample on the left and correct response (i.e., pressed any button) during catch trials as a histogram.

When using the data it might be desirable to exclude participants who had poor task performance, slept during the parts of the experiment, or experienced technical issues that resulted in missing data. Figure 5 illustrates the intersection of these various problems across participants to get a better understanding of their total impact on sample size. Data is missing for twenty-three participants due to technical errors such as button box malfunction, scanner problems, and power outages. Two participants asked to be removed from the scanner before the task was completed. A total of 50 participants have some unusable data due to poor task performance or falling asleep during the resting state or feedback scans.

**Figure 5.**
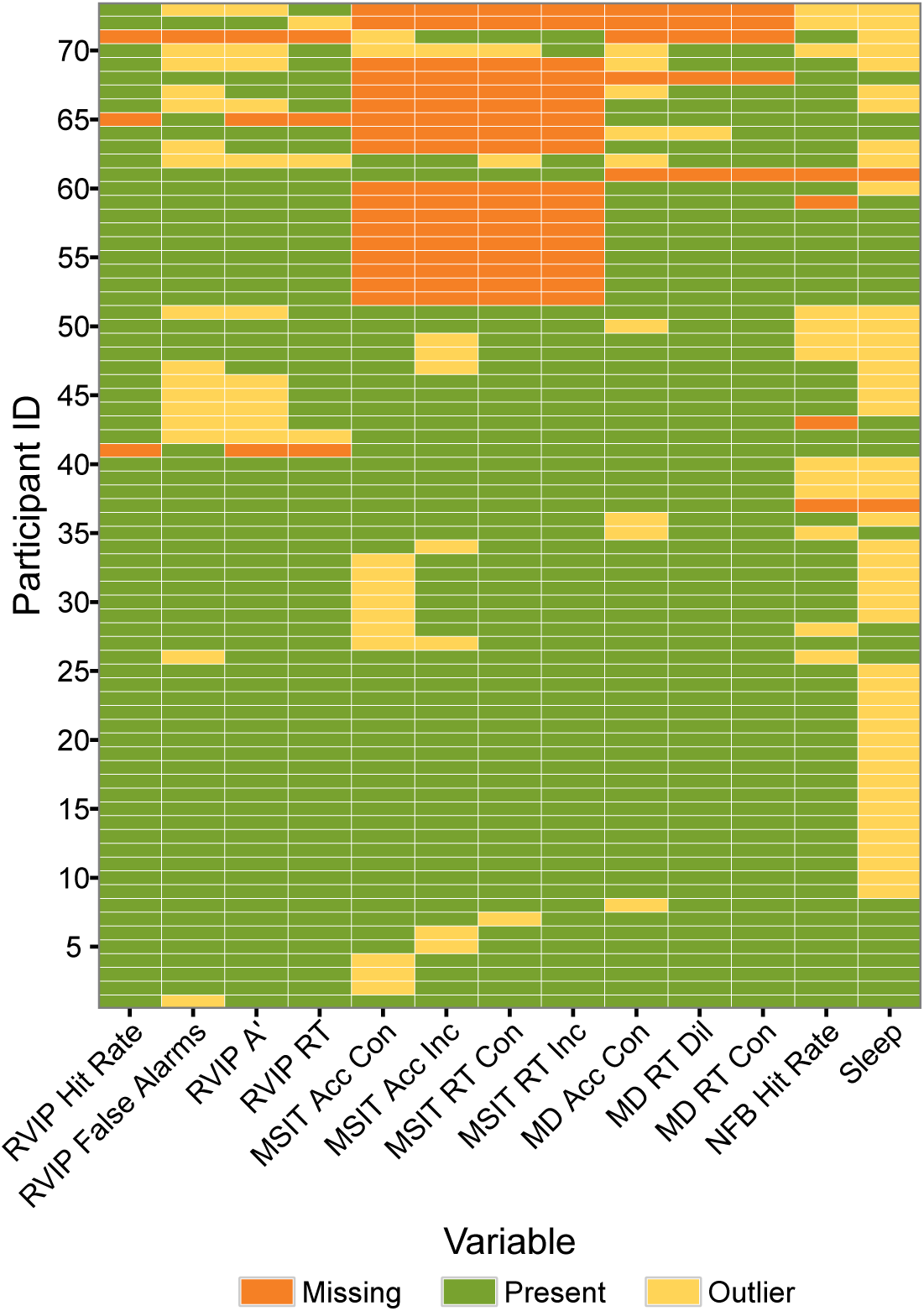
The intersection of behavioral assessment outliers and missing data. Participants are ordered by their number of outliers or missing measurements.

#### 4-3.2 Quality Assessment Protocol

To evaluate the spatial and temporal quality of the NFB fMRI data, QAP measures calculated on the data were compared to those calculated from resting state fMRI scans from the Consortium on Reproducibility and Reliability (CoRR) [77]. The CoRR dataset contains scans on 1,499 participants using a variety of test-retest experiment designs. To avoid biasing the comparison with multiple scans from the same subjects, the first resting state scan acquired on the first scanning session was used for each participant. Signal-to-noise ratio, ghost-to-signal ratio, voxel smoothness, global correlation, standardized DVARS and mean RMSD of the NFB fMRI data are all comparable to CoRR (Fig. 6). Head motion, as indexed by mean RMSD, is higher for the resting state (train) and neurofeedback scans than the other two tasks (Fig. 6F).

**Figure 6.**
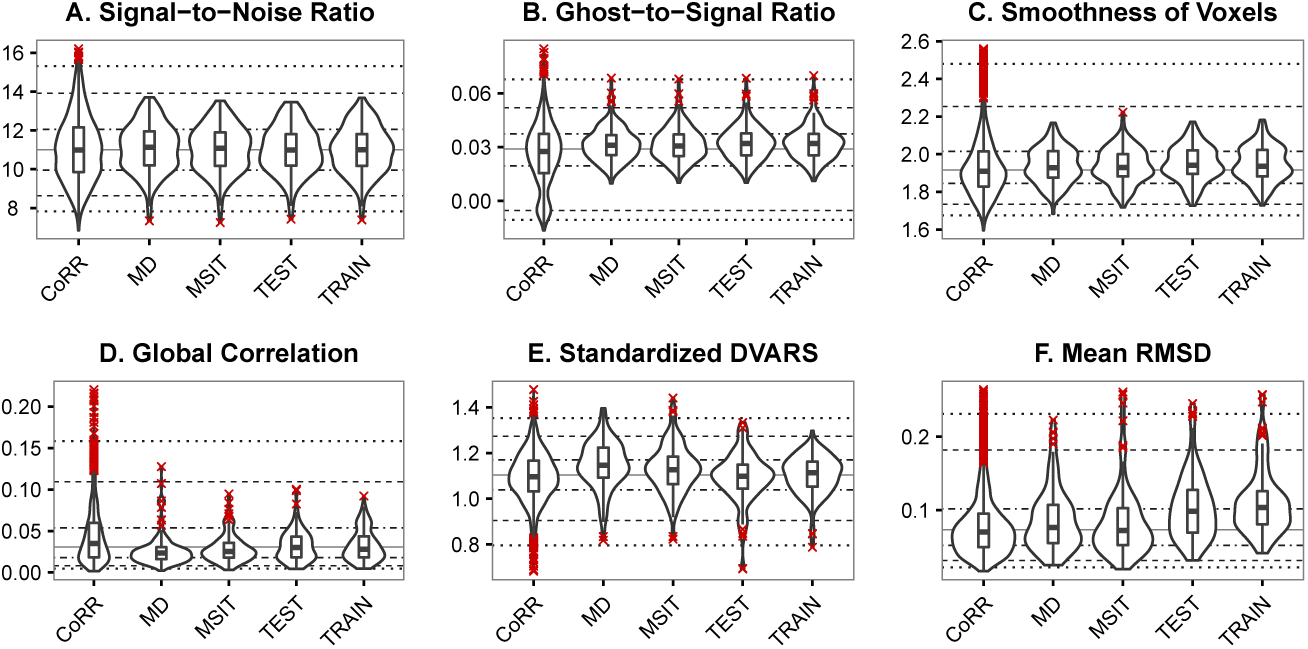
Illustrations of the overall quality of the functional neuroimaging data. Distributions of the measures for fMRI data from the MSIT, Moral Dilemma, Resting Scan (NFB Train), and Neurofeedback (NFB Test) are shown in comparison to resting state fMRI data from the CoRR dataset. Quality measures are represented as violin plots overlaid with boxplots (center line represents the median, box range represents the confidence interval, ‘X’ data points indicate outliers (1.5x Interquartile range)). Lines representing the 5^*th*^, 10^*th*^, 25^*th*^, 50^*th*^, 75^*th*^, 90^*th*^, and 95^*th*^ percentiles for the CoRR data are shown to simplify visual comparisons. Standardized DVARS is mean root-mean-square variance of the temporal derivative of voxels time courses that has been standardized to make the values comparable across different acquisition protocols. Mean RMSD is the mean root mean squared deviation (RMSD) of motion between consecutive volumes.

#### 4-3.3 fMRI Assessment

The group level analysis of the MSIT task included 87 participants after 18 were excluded for missing data and 20 were excluded for being outliers on either accuracy of response or mean reaction times (see Fig. 5). 110 participants data were included in the group level analysis of the MD task after 5 participants were excluded due to technical problems, and another 10 were excluded due to outlier task performance. As expected from the literature, the MD task robustly activated the DMN (Fig. 7A, red) and deactivated attentional networks (Fig. 7A, blue) that are typically active during task performance [8]. Consistent with previous literature [76] group level analysis of the MSIT showed robust activation of the dorsal and ventral attention networks (Fig 7B, red) and deactivation of the DMN (Fig 7B, blue). 21 participants reported falling asleep during the resting state scan and were excluded from group-level analysis. Figure 7C illustrates the expected pattern of anti-correlation between DMN and task networks [49] for the remaining 104 participants. These results confirm that these three tasks are working as expected for deactivating, activating, and localizing the DMN.

**Figure 7.**
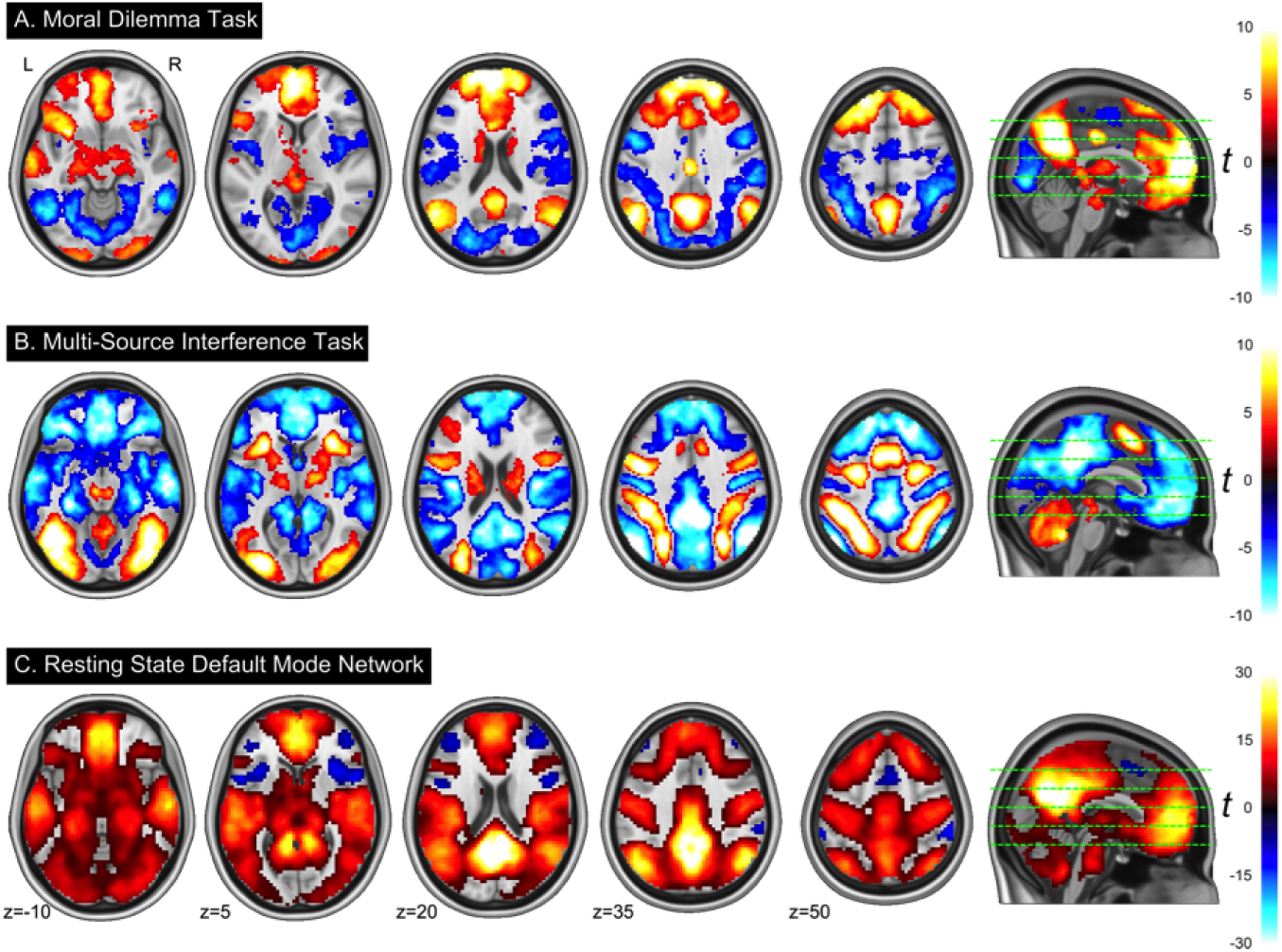
Illustrations of the overall quality of the functional neuroimagDefault network patterns extracted from the Moral Dilemma, MSIT, and Resting State fMRI Tasks. A) The dillemma > control contrast from group-level analysis of the Moral Dilemma task results in DN activation. B) The incongruent > congruent contrast of the MSIT shows DN deactivation. C) Functional connectivity of the DN extracted from the resting state task using dual regression. Statistical maps were generated by a permutation group analysis, thresholded at *p* < 0.001 TFCE FWE-corrected; overlay colors represent t statistics.

The neurofeedback task was analyzed with all of the participants that completed the scan (n=121) (Fig. 8A and B) and again with only the participants who did not fall asleep during the resting state or neurofeedback tasks (Fig. 8C). The group averaged DMN map extracted from this data is consistent with what we expect, with the exception of the prominent negative correlations (Fig. 8A). Comparing the group mean DMN time course for all participants to the task ideal time course shows that overall the task trend is followed, with a good deal of high frequency noise (Fig 8B). When the participants that fell asleep are removed, the high frequency noise remains, but the amplitude difference between wander and focus trials becomes greater, driving a higher correlation with the task waveform (Fig. 8C). This is further seen in the distribution of individual-level correlations between DMN activity and the task, where those who do not sleep perform marginally better for both conditions (Fig. 9).

**Figure 8.**
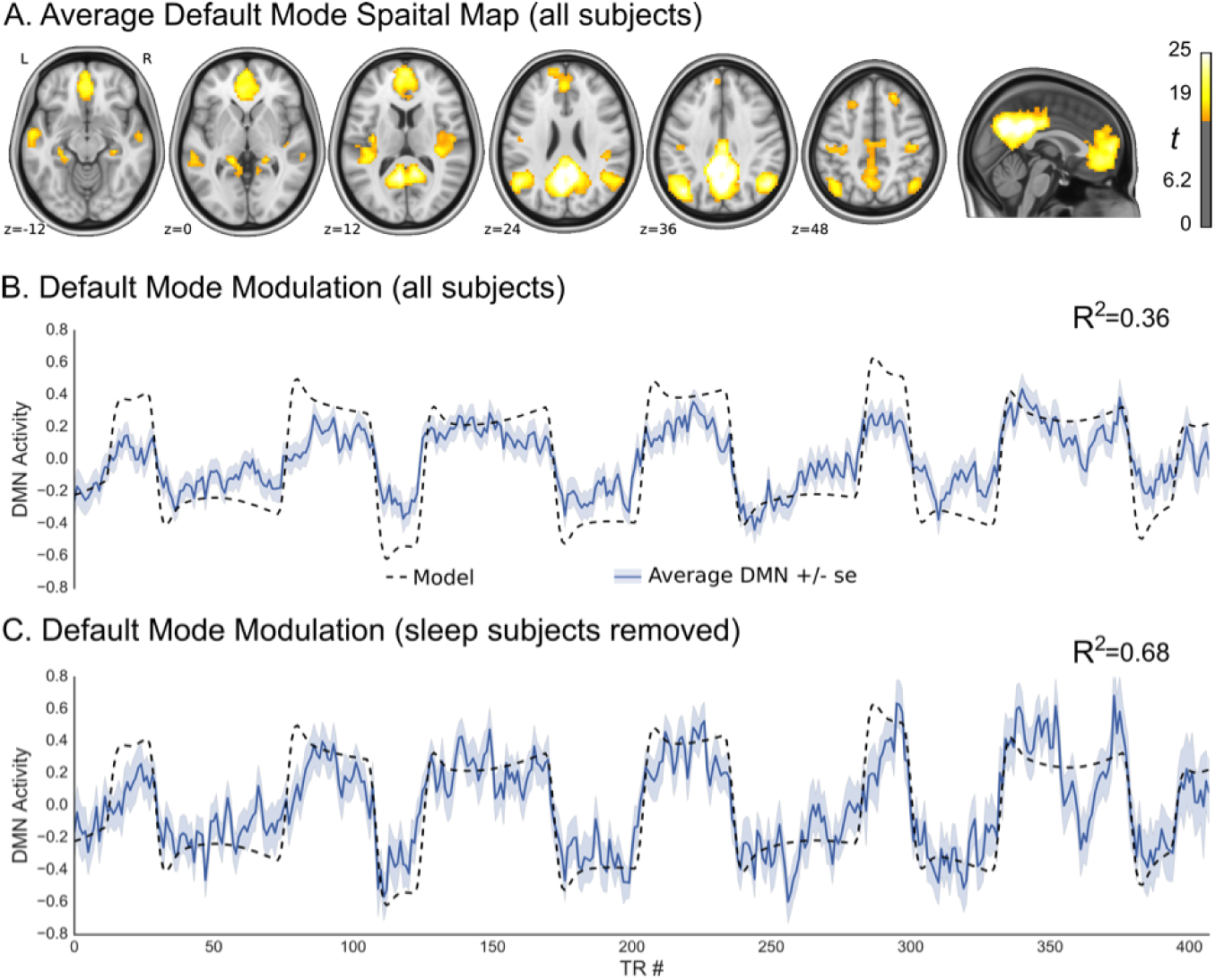
Technical validation for the neurofeedback paradigm. Panel A shows the functional connectivity map for the default mode network across all participants (p j 10-30 uncorrected, n=121) derived through dual regression. Panel B shows average overall time-series (in dark blue with shading indicating standard error) of the default mode network, across all participants (n=121), in relation to ideal time-series (in black). The coefficient of determination of the averaged time series is *R*^2^ = 0.36. Panel C shows average overall time-series (in dark blue with shading indicating standard error) of the default mode network across participants who reported no sleep during both training and feedback trials (n=76), in relation to ideal time-series (in black). The coefficient of determination of the averaged time series is *R*^2^ = 0.68.

## 5 Usage Notes

The PSWQ and PTQ were added to the assessment battery in July 2014, approximately nine months after data collection began. As a result, scores for these measures are missing from the first 26 and 27 participants, respectively. Additionally, in July of 2014, the full scale Response Styles Questionnaire (RSQ) was replaced with the newer subscale RRS, which has better psychometric properties and fewer questions. The only difference between the RRS subscale of the RSQ and the newer RRS is that one item from the Depression-Related subscale in the RRS is absent from the RSQ. To correct for this missing item in those who received the RSQ, we suggest the following: (1) Divide the RSQ derived Depression Related subscale score by 11; (2) round down to the nearest whole number (3) add this number to the RSQ derived Depression Related subscale and RSQ derived RRS-subscale total score. This procedure was validated using responses from 13 participants who received the RRS and RSQ. Correlations between scores calculated using all of the questions and those calculated using the above procedure were r = 0.9994 for the Depression-Related subscale and r = 0.9997 for the RRS total score.

For the MSIT, fMRI data should be analyzed as 42s blocks with the following onset times after dropping the first four TRs: congruent blocks - 22, 106, 190, 274; incongruent blocks −64,148, 232, 316. The Moral Dilemma fMRI data should be analyzed as 30s blocks with the following onset times after dropping the first four TRs: control blocks – 12, 72, 132, 192; dilemma blocks - 42, 102, 162, 222. Note that up until April 2014 (first 11 participants), no pre-experiment fixation period was implemented for the moral dilemma task. For those participants, all of the aforementioned stimulus onset times should be altered by subtracting 20s. For analysis of the NFB fMRI data, the first stimulus onset (and duration) times, in seconds, are as follows: 34(30), 162(60), 260(90), 418(60), 576(30), 674(90). The second stimulus onset (and duration) times, in seconds, are as follows: 68(90), 226(30), 354(60), 482(90), 610(60), 768(30). This timing information is also available in **events.tsv** files provided in the repository alongside the task fMRI data.

For some participants, the button box used to record participant responses in the scanner was defective, resulting in unusable data for the MSIT task (18 participants) and Moral Dilemma task (5 participants).

As previously discussed scripts are provided with each of the tasks for calculating response accuracies and reaction times. The assumptions made in calculating these scores may not be appropriate to all researchers, or in all applications. For this reason we have released the trial-by-trail response information in log files that accompany the fMRI data.

The impact of sleep on intrinsic brain activity, and the preponderance of sleep that occurs during resting state fMRI acquisition, has been highlighted in the literature [78, 79]. Indeed, a large proportion of the participants in this study reported falling asleep during either the resting state or neurofeedback scans. Whether or not this data should be excluded is up to the researcher and depends on the analysis being performed. Users of this resource might also consider whether they trust the self reports, or whether they should try to decode an objective measure of sleep from the data [79]. Additionally, researchers could potentially use the respiratory, heart rate, or galvanic skin response recordings provided in the repository to monitor wakefulness.

See information in Data Privacy section of Data Records for restrictions and limitations on data use.

## 6 Discussion/Conclusions

This manuscript describes a repository of data from an experiment designed to evaluate DMN function across a variety of clinical and subclinical symptoms in a community-ascertained sample of 180 adults (50% females, aged 21-45). The data includes assessments that cover a variety of domains that have been associated with or affected by DMN activity, including emotion regulation, mind wandering, rumination, and sustained attention. Functional MRI task data is included for tasks shown to activate, deactivate, and localize the DMN, along with a novel real-time fMRI neurofeedback paradigm for evaluating DMN regulation. Preliminary analysis of the first 125 participants released for sharing confirms that each of the tasks is operating as expected.

Group level analysis of the neurofeedback data indicates that the participants are able to modulate their DMN along with the task instructions. For all participants the group average time-course co-varies significantly with the task model and increases when removing participants who reported sleep. Greater than half of the participants who reported no sleep had a significant correlation (p <0.05; r>0.15, phase randomization permutation test) with the task model (see Fig. 9). The large portion of participants who were able to significantly modulate their DMN shows the effectiveness of the neurofeedback protocol.

**Figure 9.**
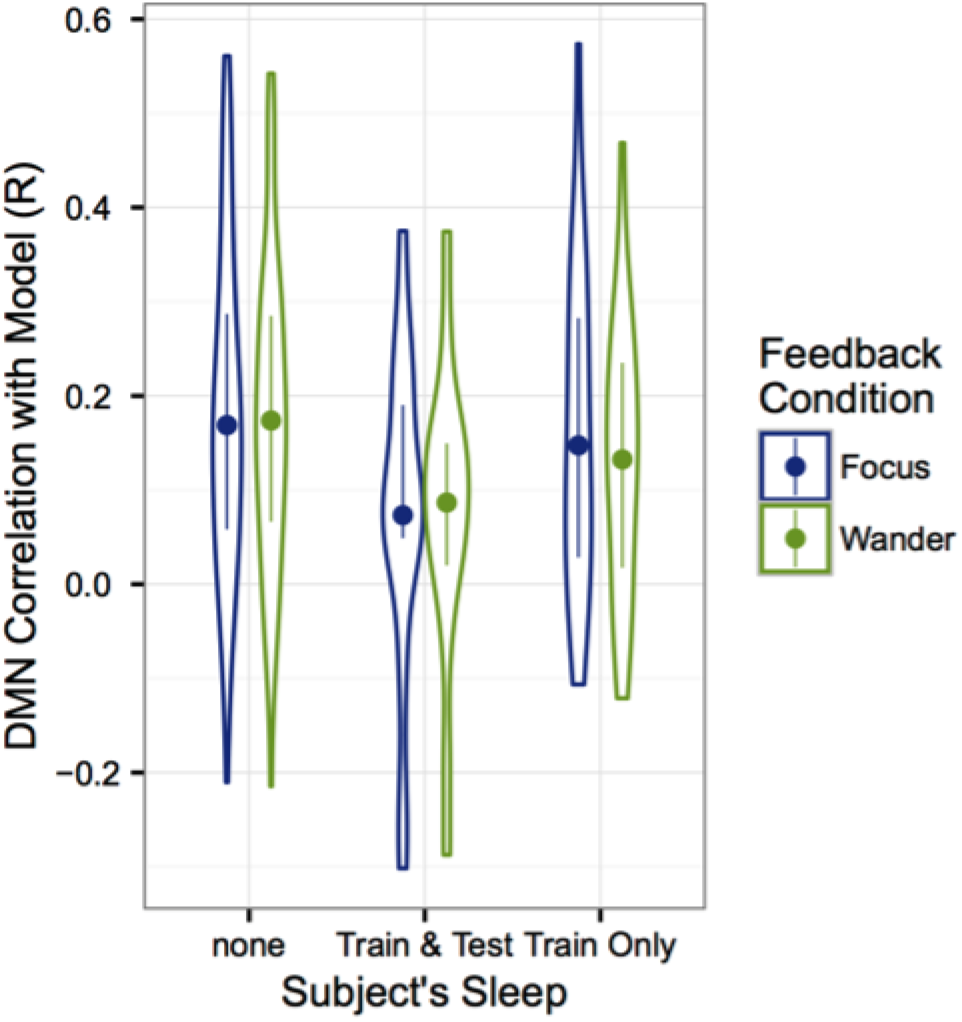
Distribution of individual correlations for participant default mode network time-series with ideal-times series as a function of sleep status. Each sleep status group is then sub-divided into the two task conditions – Focus/Wander. Subjects with reported sleep during both trials show lower correlations with the model compared to subjects with no reported sleep. The dots indicate the mean within each group and the distributions of the correlations are plotted around the mean.

One unexpected result of our technical validation is a high amount of data loss due to poor participant performance and compliance. Excluding all of the participants who are considered outliers on at least one of the tasks or who fell asleep during the resting state or neurofeedback scans, would remove 73 of the 125 participants (see Fig. 5). Forty-six individuals could be excluded for sleep, which is not surprising given recent reports of the high incidence of sleep during resting state scans [79]. Interestingly, 26 of these participants were also outliers in at least one other task. This indicates that either they had trouble staying awake during the other tasks as well, or were non-compliant overall. Due to a paucity of information on participant compliance during fMRI experiments, it is hard to tell whether what we are seeing is common for our population, or whether there is a specific problem with our experiment protocol. We do believe that compliance would be higher if we were to utilize a younger healthy population, or participants who have been scanned multiple times, as is commonly done in cognitive neuroscience. These problems with sleep and poor performance illustrate the need to debrief participants, monitor their wakefulness during the scan, or try decode sleep from the fMRI data [79]. Additionally, it is important to check the data as it is acquired so that a study protocol can be adapted to reduce data loss.

Although much of the interest in real-time fMRI based neurofeedback is focused on clinical interventions [80], it is also a valuable paradigm for probing typical and atypical brain function. We believe that the data in this repository will have a substantial impact for understanding the nuances of DMN function and how variation in its regulation leads to variation in phenotype. This resource will be particularly useful to students and junior researchers for testing their hypothesis of DMN function, learning new techniques, developing analytical methods, and as pilot data for obtaining grants. We encourage users to provide us with feedback for improving the resource and are very interested to learn about the research performed with the data.

## Author’s contributions

RCC, MPM, SJC, FXC and BL designed the experiment, RCC and SLC developed the real-time DMN tracking system, SLC and JL provided the custom real-time fMRI sequence, ARM, NTVD, JM, and RCC wrote the manuscript, ARM, JM, CF, SG and RCC performed data analyses, JP, ARM, BP, and CF organized data for release, RCC, CCCB, ARM, NTVD, and MB designed and implemented the behavioral tasks and assessments, RT, AMB, RCC, MPM and SJC managed data collection, AA, MMB, ARM, SC, TPC, CG, AG, JG, SH, MK, AL, AMB, LP, HR, CS, ES, RS, MS, ET, KT, BV, LW, and ABW were involved in participant recruitment and performed data collection, VDC, MK, DL, BM, RW, and DW managed data sharing through the COINs pipeline.

## Acknowledgements

We would like to thank Cathy Hu and Raj Sangoi for MRI operation. Data collection and salary support were provided by NIMH BRAINS R01MH101555.

